# Dynamic reorganization of brain functional networks during cognition

**DOI:** 10.1101/012922

**Authors:** Michał Bola, Bernhard A. Sabel

**Affiliations:** Institute of Medical Psychology, Medical Faculty, Otto-von-Guericke University of Magdeburg, Magdeburg, Germany.

**Keywords:** Brain networks, graph theory, functional connectivity, “the connectome”, perception, cognition

## Abstract

How cognition emerges from neural dynamics? The dominant hypothesis states that interactions among distributed brain regions through phase synchronization give basis for cognitive processing. Such phase-synchronized networks are transient and dynamic, established on the timescale of milliseconds in order to perform specific cognitive operations. But unlike resting-state networks, the complex organization of transient cognitive networks is typically not characterized within the graph theory framework. Thus, it is not known whether cognitive processing merely changes strength of functional connections or, conversely, requires qualitatively new topological arrangements of functional networks. To address this question, we recorded high-density EEG when subjects performed a visual discrimination task and conducted and event-related network analysis (ERNA) where source-space weighted functional networks were characterized with graph measures. We revealed rapid, transient, and frequency-specific reorganization of the network’s topology during cognition. Specifically, cognitive networks were characterized by strong clustering, low modularity, and strong interactions between hub-nodes. Our findings suggest that dense and clustered connectivity between the hub nodes belonging to different modules is the “network fingerprint” of cognition. Such reorganization patterns might facilitate global integration of information and provide a substrate for a “global workspace” necessary for cognition and consciousness to occur. Thus, characterizing topology of the event-related networks opens new vistas to interpret cognitive dynamics in the broader conceptual framework of graph theory.

## 1. Introduction

Studying the brain “connectome” is an interdisciplinary and rapidly developing field with the premise that understanding organization of brain networks will lead to a major leap forward in basic and clinical neuroscience. Brain connectivity networks exhibit complex structures and emergent topological properties which cannot be reduced to just the sum of pairwise interactions. Rather these topological features constitute a whole new level of brain analysis. Therefore, patterns of anatomical and functional connectivity have been extensively studied with graph theory measures (review: Sporns et al., 2004; Bassett and Bullmore, 2006; Stam, 2010; Park and Friston, 2013).

In graph theory topological arrangements are thought to facilitate or hamper the network’s information processing capabilities (Newman, 2003; Sporns, 2013). Neuroscience provides support for this hypothesis as, indeed, topology of brain resting-state functional networks predicts individual cognitive performance (van den Heuvel et al., 2009; Langer et al., 2012). Furthermore, functional networks reorganize towards more advantageous arrangements during development (Boersma et al., 2011) but shift back towards a less optimal structure in aging (Wang et al., 2010). Disrupted network organization is also a hallmark of numerous pathological conditions including Alzheimer’s disease (Stam et al., 2009), traumatic brain injury (Caeyenberghs et al., 2012), unipolar depression (Lord et al., 2012), and vision loss (Bola et al., 2014).

Dynamics of brain network reorganization can be studied on multiple time-scales (Kopell et al., 2014). By studying how steady, long-lasting changes in topology of anatomical and functional networks are related to changes in cognitive capabilities we have greatly advanced our understanding of neural development and neurodegeneration. But transient cognitive networks, established and dissolved on the timescale of milliseconds, are typically not characterized in terms of their topological arrangements. It is not clear whether networks’ structure reorganizes during cognitive processing and, if it does, what is the specific reorganization pattern. Two scenarios might be considered: Firstly, cognitive operations might merely change the weights of connections with networks’ global topology remaining fixed, not modifiable on the fast time-scale. Several neurophysiological (Bassett et al., 2006; Nicol et al., 2012; Betti et al., 2012; Jin et al., 2012) and fMRI studies (Cole et al., 2014) lend support to this hypothesis. The second possibility is that cognition might involve qualitatively new topological arrangements of networks. Such reorganization was postulated by the global workspace theory (Baars, 2002; Baars et al., 2013; Dehaene and Changeux, 2011) and supported by a number of MEG/EEG (Valencia et al., 2008; Doron et al., 2012; Palva et al., 2010; Kitzbichler et al., 2011; de Vico Fallani et al., 2008) and fMRI studies (Bassett et al., 2011, 2013; Ekman et al., 2012; Mennes et al., 2013).

Discovering the pattern of cognition-related reorganization would open new vistas to interpret cognitive operations within the conceptual framework of graph theory and network science. We therefore set out to study topological properties of brain functional networks during visual perception and cognition. In the present study we employed a classic visual oddball task to probe cognitive processing. In this task identification and processing of infrequent ‘oddball‘ targets involves several cognitive operations and activates distributed parieto-frontal regions (Brázdil et al., 2007; Kim, 2014; review: Polich, 2007). Because the oddball task is commonly used as a model of cognitive processing, here it was employed to study network reorganization. We recorded high-density EEG during task performance and conducted an event-related network analysis (ERNA) where topology of source-space phase-synchronized networks was characterized with graph measures on a millisecond timescale. We hypothesized that cognitive processing in the oddball task is related to rapid, transient, and frequency-specific topological reorganization of brain functional networks.

## 2. Methods

### 2.1 Subjects

We tested 18 right-handed subjects without any neurological or neuropsychiatric disorders, with normal or corrected-to-normal visual acuity. All subjects gave informed consent before the experiment. Data of 2 subjects were discarded due to poor EEG quality (i.e. excessive number of electrodes recording artefacts due to poor contact with the scalp); thus data of 16 subjects (8 female; 25±0.7 years old) were included in the analysis.

### 2.2 Experimental setting

Participants performed a classic visual oddball task. The experiment comprised two conditions: presentation of frequent distractors (DIST; 1180 trials; 90% probability) and rare targets (TARG; 120 trials; 10% probability). Subjects were instructed to ignore distractors and to only press a space key on a keyboard as fast as possible in response to targets. Gabor patches, created by multiplying a 2D Gaussian (SD=2deg. of visual angle) and 2D sinusoidal grating (contrast=70%, spatial frequency=1cycle/deg), were used as stimuli. They were presented in the center of the screen and orientation of a patch (vertical vs. horizontal) distinguished distractors from targets. The orientation of targets was counterbalanced across subjects. Subjects viewed stimuli binocularly from the distance of 57cm on a gamma corrected monitor (EIZO, CG241W). In total, 1200 stimuli presentations were given, grouped into 4 blocks of 300 stimuli, with short breaks in between. A red fixation point (size 0.25deg) was present on the screen all the time except for the stimulus presentation and the subject was asked not to move the eyes away from it. The stimuli were presented for 200ms and the maximum allowable response time was 1200ms. After the response time a random inter-trial interval was started (range 200-800ms). Each subject performed a few practice trials before commencing the data collection.

### 2.3 EEG acquisition

Dense array EEG was recorded using a HydroCell GSN 128-channel net and Net Amps 300 amplifier (EGI Inc., Eugene, Oregon, USA). The signal was sampled with 500Hz frequency, referenced to Cz (ground electrode between Cz and Pz), and digitalized with 24 bit precision. Impedance was ascertained to be below 100kΩ throughout the recording.

### 2.4 EEG preprocessing

Continuous EEG signal was re-referenced to the linked mastoids, filtered with high-pass (1Hz) finite impulse response (FIR) filter, low-pass (100Hz) FIR filter, notch (50Hz) FIR filter, and down-sampled to 250Hz. The EEG signal was divided into epochs time locked to the stimuli [-0.8s to 1.7s] which were baseline [-0.2s to 0s] corrected. Epochs were screened for non-stereotypical artefacts e.g. excessive myographic activity (on average 50±9 epochs per subject discarded). Noisy channels were discarded (6±1.9 channels per subject discarded) and interpolated based on the activity of surrounding channels. Independent component analysis (ICA) was performed (Bell and Sejnowski, 1995). Topographic maps, power spectra, and time-domain activity was evaluated visually for each component and based on these features each component was classified as representing either brain activity or artefacts e.g. eyeblinks or cardiac activity. Components representing artefacts were discarded. On average 15.8±1.3 components were retained and back-projected into sensor space.

Target trials not followed by a response (omissions), and distractor trials followed by a response (false positives) were discarded. To exclude any bias from the unequal number of trials in both experimental conditions only a subset of distractor trials, for each subject equal to the number of target trials, was randomly chosen for further analysis. On average 117±1 trials per subject/condition were analyzed.

The frequency bands were defined as follows: theta (4-7Hz), alpha (7-14Hz), beta (14-30Hz).

### 2.5 Source-reconstruction procedure

The forward model and the inverse model were calculated with an open access software Brainstorm (Tadel et al., 2011). The forward model, which describes the signal pattern generated by a unit dipole at each allowed location on the surface, was calculated using the symmetric boundary element method (BEM) (Gramfort et al., 2010, Kybic et al., 2005) and default MNI MRI template (Colin 27). Preprocessed, ICA pruned, stimulus-locked single-trial data were used to calculate the inverse model, which was estimated using the weighted Minimum Norm Estimate (wMNE) (Hämäläinen and Ilmoniemi, 1994). wMNE is well-suited for estimation of large-scale functional connectivity networks as it addresses the volume conduction and thus reduces spurious signal correlations (Hassan et al., 2014; Palva and Palva, 2012). When computing the inverse operator (a) the source orientations were constrained to be normal to the cortical surface; (b) a depth weighting algorithm was used to compensate for any bias affecting the superficial sources calculation (Lin et al., 2006); and (c) a regularization parameter, *λ*^2^ = 0.1 was used to minimize numerical instability, reduce the sensitivity of the wMNE to noise, and to effectively obtain a spatially smoothed solution (Hämäläinen and Ilmoniemi, 1994). In this way, activation time-courses at 15002 vertices (an equilateral triangle in the tessellation of the cortical surface) were estimated. The cortical surface was divided into 68 anatomical regions of interest (ROI; 34 in each hemisphere) based on the Desikan–Killiany atlas (Desikan et al., 2006) and activity of a seed voxel of each area was used to calculate functional connectivity.

### 2.6 Spectral decomposition

The main steps of the conducted analysis are presented in **Figure 1**. Spectral decomposition of EEG from single trials (both, sensor-space and source-space) was conducted with Morlet wavelet (EEGlab *newtimef* function) and 200 linearly spaced (every 8ms) time points and 40 linearly spaced (every ≈0.7Hz) frequencies were estimated. The window size used for decomposition was 211 data points (844ms), thus the overlap between windows was ≈99% . The wavelet contained 3 cycles at the lowest frequency (3.9Hz) and the number of cycles was increasing up to 11.4 cycles at highest frequency (30Hz) and 40 frequency points linearly spaced between 3.9Hz and 30Hz were estimated. Therefore, for every subject, condition, and channel/ROI, we obtained a 3D matrix of 40 (frequency points) X 200 (time points) X ‘number of trials’ (which varied between participants). Absolute values of the decomposed sensor-space signals were analyzed to investigate event-related changes in oscillatory power.

**Fig 1.**
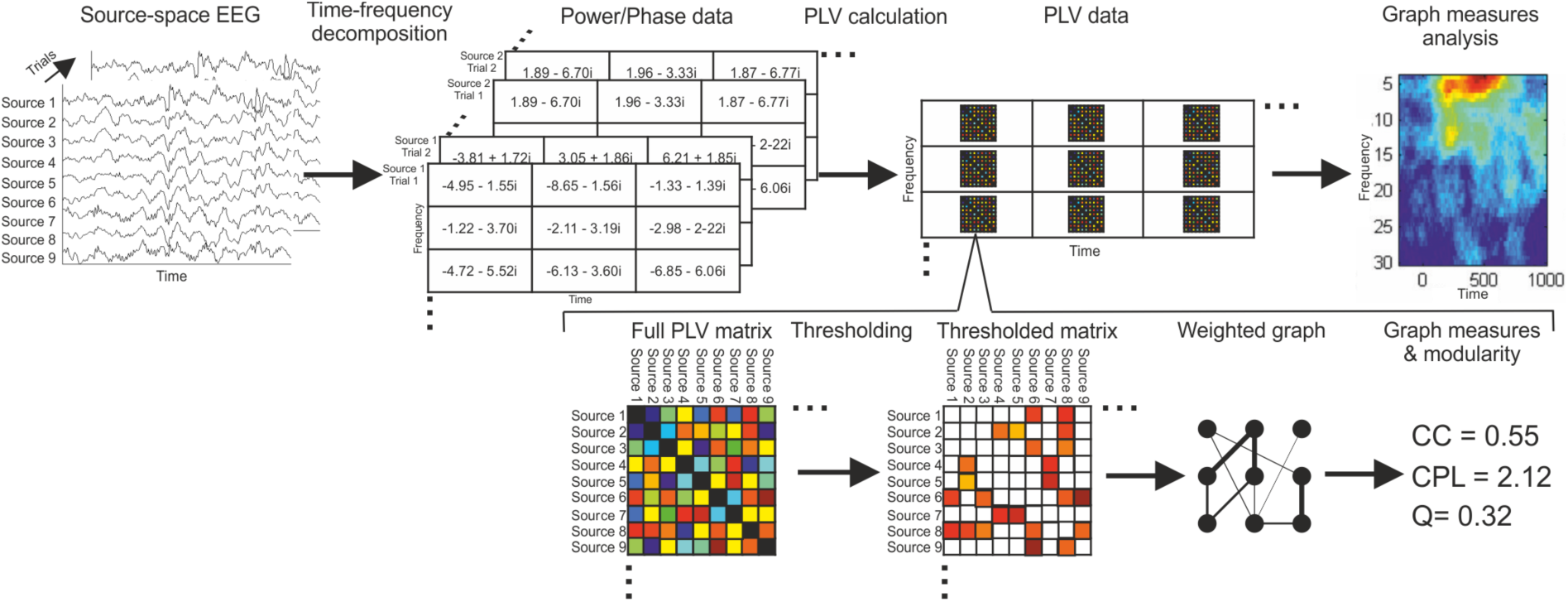
Analysis pipeline. Data are preprocessed, divided into stimulus-locked epochs, and projected into the source-space using the weighted MNE algorithm. Signals of the seed voxels of 68 anatomical brain regions are decomposed with a Morlet wavelet. For each time- and frequency-point a full adjacency matrix containing *PLV* estimates is created (in the figure color represents coupling strength). Each *PLV* matrix is thresholded to create sparse, weighted, undirected graph. Topology of each graph is characterized with graph measures. Graph measures are then represented in the time-frequency space and tested statistically.

### 2.7 Functional connectivity estimation

Two measures of coupling were calculated for each frequency- and time-point between all pairs of ROIs. Phase Locking Value (*PLV*; Lachaux et al., 1999), which measures variability of phase between two signals across trials, is classically defined as,

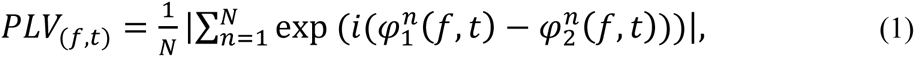

where 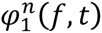 and 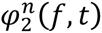 denotes phase from ROI 1 and 2 respectively, from trial *n* and for frequency-point *f* and time-point *t*. *N* denotes the number of trials, *i* is the imaginary unit, and |*x*| indicates an absolute value of *x*.

Imaginary part of coherence (*iCoh*; Nolte et al., 2004), which is a conservative measure of functional coupling insensitive to volume conduction, was calculated as,

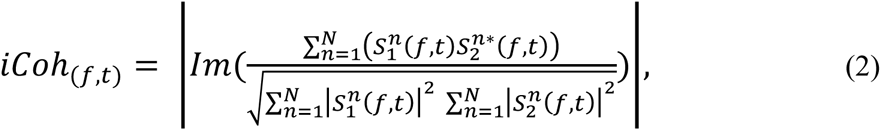

where 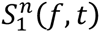 and 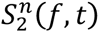 are wavelet-decomposed EEG signals from ROIs 1 and 2 respectively, * indicates the complex conjugate, and |*x*| indicates an absolute value of *x.* For every subject, condition, and all pairs of ROIs (2278 pairs) we obtained *PLV* and *iCoh* matrices 40 (frequency points) X 200 (time points). In other words, for every subject (n=16), condition (2), and time- (200), and frequency point (40) we obtained a full 68 X 68 adjacency matrix.

### 2.8 Weighted graphs

The main aim of the study was to analyze topology of the event-related functional networks. To this end we converted full *PLV* adjacency matrices into sparse, undirected, weighted graphs which can be analyzed with graph measures. Graphs comprise of nodes, being systems’ elements (here brain areas), and edges/connections, indicating interactions between elements (here phase synchronization). In order to obtain a sparse, weighted, undirected graph/network ***A****_(f,t)_*, a full adjacency matrices were thresholded, so that all the values below the threshold were set to 0. The values above the threshold retained original values (weights). For each matrix the threshold was individually adjusted, so that the *density*, defined as the proportion of existing edges out of all possible edges, was equal for each graph. The main analysis was conducted with networks of *density*=0.29. Yet, the control analyses indicated that the main effects found in the study are independent of networks’ *density*.

The basic parameter which can be calculated for weighted networks is networks’ *strength* defined as an average over weights of all connections within a network. Further *nodal strength* can be calculated for each node defined as sum of weights of all edges coupled to a node.

### 2.9 Graph measures

#### 2.9.1 Weighted networks

Weighted graphs ***A****_(f,t)_* were characterized with several graph measures generalized for analysis of weighted networks as implemented in Brain Connectivity Toolbox. Formal definitions of all measures can be found in (Rubinov and Sporns, 2010). Clustering Coefficient (*CC_(f,t)_*) and Characteristic Path Length (*CPL_(f,t)_*) were calculated with functions *clustering_coef_wu* and *charpath/distance_wei*, respectively.

Further, we calculated the weighted Rich Club Coefficient (*RCC*). Many systems exhibit so called rich-club organization, meaning that nodes with high *degree* (i.e. network hubs) are more strongly interconnected among themselves than nodes of a low degree (Colizza et al., 2006). The anatomical networks of the human brain were shown to exhibit the rich-club topology (van den Heuvel and Sporns, 2011). *RCC* quantifies how strong the interactions among networks’ hubs are. To calculate *RCC* all edges of the analyzed graphs were ranked by weight, resulting in a vector *W^ranked^*. *RCC* is typically calculated for a range of *rcK* values and for each value of *rcK*, a group of nodes with *degree*>*rcK* belongs to a rich-club. The number of edges *E_rc_* between the rich-club nodes was counted, together with their collective weight *W_rc_* calculated as the sum of weights of all rich-club edges. The weighted *RCC* was then calculated as the ratio between *W_rc_* and the sum of weights of the strongest *E_rc_* edges of the whole graph, given by the top *E_rc_* number of edges of the collection of ranked edges in *W^ranked^*. *RCC* was formally was defined as follows (Opsahl et al., 2008):
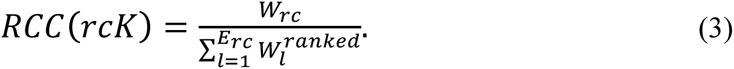

The main analysis was conducted with *rcK*=25 but all the individual rich-club curves are also presented. *RCC* was calculated with the Brain Connectivity Toolbox function *rich_club_wu.*

Finally, k-core decomposition of graphs was calculated with a *kcore_bu* function. For this analysis weighted graphs were binarized, as otherwise the network *strength* would affect results to a great extent. K-core decomposition defines maximal connected sub-graphs in which all nodes have *degree*>*k*. K-core decomposition is implemented in steps. In each step nodes (together with their edges) with *degree*<*k* are pruned. Then *k* is increased and again the nodes with *degree*<*k* are pruned, until all nodes are removed. For each step removed nodes have the value *k* assigned as the *nodal k-core* value. A high *nodal k-core* indicates a more central role of a node in a network. The subset of nodes pruned in the last step (i.e. set of nodes with highest *nodal k-core*) constitute the k-core of a network.

#### 2.9.2 Binary graphs

The same procedures were followed to create binary graphs. Full adjacency matrices were thresholded to obtain graphs with the same *density* (0.29) as the weighted graphs, but now all the preserved entries (i.e. above the threshold) were set to 1. To analyze binary graphs the same graph measures (but implemented specifically for analysis of binary graphs) were used. Clustering coefficient and characteristic path length were calculated with functions *clustering_coef_bu* and *charpath/distance_bin* respectively. Binary *RCC* was calculated with the *rich_club_bu* function.

### 2.10 Modularity

A module (community) is defined as a group of nodes which are more strongly connected among each other than with nodes in other modules. Typically a quality function *Q* is used to evaluate modularity of a network (Newman and Girvan, 2004). In binary networks *Q* quantifies the number of intra-modular connections relative to the inter-modular connections. In weighed networks *Q* quantifies the weights of intra-modular connections relative to the inter-modular connections. Therefore, maximization of *Q* allows partitioning a network into the most optimal community structure.

#### 2.10.1 Uni-layer and multi-layer networks

We firstly studied modularity of uni-layer weighted networks. Each network ***A****_(f,t)_* represents one time- and frequency-point thus modularity at each time-point was evaluated independently from modularity at other time-points. ***A****_(f,t)_* is a 68 X 68 matrix whose elements ***A****_ij_* specific weights of edge between nodes *i* and *j*. Suppose that node *i* is assigned to community *g_i_* and node *j* is assigned to community *g_j_*. The quality function *Q_uni_* evaluating modularity of a uni-layer network can be defined as:

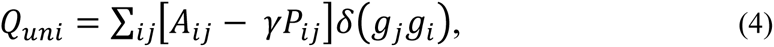

where *δ* is the Kronecker delta, and thus *δ*(*g*_*j*_*g*_*i*_) = 1 if *g*_*j*_ = *g*_*i*_ and 0 otherwise, and *γ*(*gamma*) is a resolution parameter. *P*_*ij*_ is the expected weight of the edge connecting node *i* and node *j* under the Newman-Girvan null model, which was defined as,
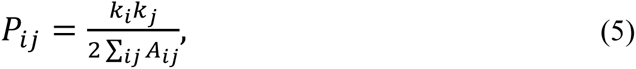

where *k_i_* is the strength of node *i* and *k_j_* is the strength of node *j*. The resolution parameter *γ* was set to 1 in the main analysis but we repeated the analysis with different values of *gamma* to demonstrate robustness of the results.

Further, for each frequency we created multilayer networks ***B****_(f)_* where each layer (“slice”) of a network represented networks’ state at one time point. Each layer was linked to proceeding (t-1) and subsequent (t+1) layers. The temporal links allowed estimating evolution of the community structure in the time domain. Neuroimaging data have been already studied as multilayer networks (Basset et al., 2011, 2013; Doron et al., 2012). To ensure that mainly the evoked component is represented, and not the pre-stimulus activity, only the time-points from 0ms to 800ms after the stimulus onset were included in the multi-layer networks. Thus each multilayer network comprised 96 layers (time-points). The quality function *Q_ml_* evaluating modularity of a multilayer network can be defined as,

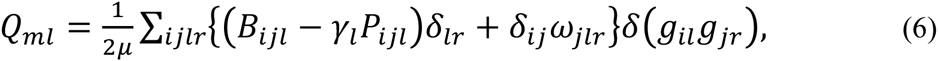

where *δ* is the Kronecker delta, the adjacency matrix of layer *l* has components *B_ijl_*, the element *P_ijl_* gives the components of the corresponding null model matrix, and *γ_l_* is the structural resolution parameter of layer *l*, *g_il_* gives the community assignment of node *i* in layer *l*, *g_jr_*, gives community assignment of node *j* in layer *r*, the parameter *ω*_*jlr*_ (*omega*) is the interlayer coupling strength between node *j* in layer *r* and node *j* in layer *l*, 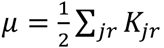, the strength of node *j* in layer *l* is *K_jl_* = *k_jl_* + *c_jl_*, the intra-layer strength of node *j* in layer *l* is *k_jl_*, and the inter-layer strength of node j in layer l is 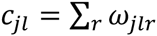. The Newman-Girvan null model was employed within each layer and defined as,
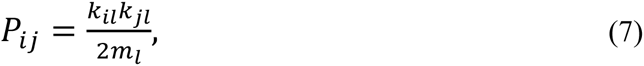

where 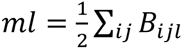 is the total edge weight in layer l. The resolution parameter *γ_l_*(*gamma*) was set to 1 and all non-zero *ω_jlr_* were set to 1 in the main analysis. The multilayer analysis was repeated with different values of *gamma* and *omega* to demonstrate robustness of results.

#### 2.10.2 Analysis of networks partitions

We used a Louvain greedy community detection algorithm (Mucha et al., 2010) to optimize *Q_uni_* and *Q_ml_*. Networks were partitioned into non-overlapping communities, i.e. each node belonged to one community only. Importantly, algorithms partitioning networks by optimization of modularity function tend to produce many near-optimal partitions which form the so called high-modularity plateau. Hence, two partitions optimizing the function Q to the same degree might indicate different community structure. To deal with this problem the high-modularity plateau is typically extensively sampled, i.e. that same network is partitioned a number of times. Having several degenerate partitions of the same network there are two possibilities: (i) a single “representative” partition might be chosen and analyzed (e.g. Doron et al., 2012); (ii) measures of interest (e.g. number of modules) might be calculated from every obtained partition and then averaged over all partitions to give a representative value for the underlying network (e.g. Bassett et al., 2011). Here we chose the latter approach. Thus, each uni-layer and multi-layer network was partitioned 25 times, features of interest were calculated from each partition and averaged over all 25 degenerate partitions to obtain the representative value for the network. The averaged features were then compared statistically.

The features of partitions analyzed in the uni-layer networks were: (i) *Q_uni_* and (ii) number of modules. The number of modules was calculated separately for each time point. The features of partitions analyzed in multilayer networks were: (i) *Q_ml_*, (ii) number of modules, and (iii) flexibility. Here the number of modules reflected the number of modules throughout the temporal network. Flexibility was defined as number of the times a node changes community assignment divided the number of all possible changes (Bassett et al., 2011).

### 2.10.3 Null network models

Importantly, investigating qualities of the community structure is of relevance only if the analyzed network is indeed modular. Therefore, before comparing modularity between experimental conditions we tested whether event-related networks exhibit modular structure by comparing them to null (surrogate) networks. Null networks were created by destroying the possible modular structure of original networks by randomization.

For each uni-layer network ***A***_(t,f)_ we created 25 null networks by randomizing edges of the original network but preserving weight, degree, and strength distributions (function: *null_model_und_si*). See Rubinov and Sporns (2011) for details of the algorithm

When analyzing multilayer networks we used three different null models proposed by Bassett et al., (2011): (i) we randomized edges in each layer, but preserved weight, degree, and strength distributions (function: *null_model_und_si*; the same procedure as for static networks); (ii) we randomized nodes in each slice so that node A in slice *t* was not linked to node A in slice *t*+1 but to some other node; (iii) we randomized slices, so that slice *t*+1 did not follow slice *t*. For each multilayer network ***B***_(f)_ we created 25 instantiations of each null model. Each null network was partitioned into communities using the same procedure as for the original networks. From each partition network features were calculated and averaged over 25 null networks. Difference between features of original networks and null networks was calculated and tested against 0.

Due to high computational time needed to calculate numerous network partitions we limited the community structure analysis to three frequencies representing 3 frequency bands (3.9Hz – theta; 10.6Hz – alpha; and 20.6Hz – beta).

Analysis of community structure in binary graphs was conducted according to the same procedures.

### 2.11 Control analyses

When analyzing topology of sparse networks one has to define a number of parameters which might influence the result of the analysis. The most important parameter is *density* of analyzed graphs. When estimating modularity a resolution parameter *gamma* has to be defined and when estimating multi-layer modularity also a between-layer coupling also a parameter *omega* needs to be set. Thus, to demonstrate the main results were robust against varying these three parameters we conducted a number of control analyses. We focused on the main effects found between conditions which were: (i) reorganization of the theta-band network around 350-550ms shown by *CPL* and *CC*; (ii) decrease of modularity of the theta network around 350-500ms and beta network around 600-700ms in uni-layer networks. Alpha network 300-500ms was also included in the control analysis although there was no effect in the alpha band; (iii) reorganization of theta and beta multi-layer networks (alpha network also included in the control analyses). For each value of the parameter the difference (Δ) between conditions (i.e. TARG-DIST) calculated for each subject and tested statistically against 0.

### 2.12 Statistical analysis

In the present study we sought to characterize event-related changes in network topology. Thus, the EEG measures (*PLV*/*iCoh*, graph measures) were compared: (i) against baseline which was defined as time window [-200ms – 0ms] with respect to the stimulus onset (measures of the “baseline state” were calculated for each frequency separately); and (ii) between conditions (DIST vs. TARG). The comparisons involving multiple time-points (ERP) or multiple time-frequency points (*PLV*/*iCoh*, graph measures) were conducted using repeated measures cluster mass permutation test to deal with the problem of multiple comparisons (Bullmore et al., 1999; Maris and Oostenveld, 2007).

When studying networks modularity we additionally conducted comparisons between original networks and surrogate (random) networks. Here for each subject a difference (REAL-SURROGATE) was calculate and tested against zero with a two-tailed repeated-measures t-test. The difference between conditions was considered significant when *p*<0.05

Cluster mass permutation test allows controlling for the false positive rate when multiple comparisons are conducted. It is especially useful when there is no *a priori* hypothesis as to when, where, and in which frequencies the effect might occur. When testing ERP time points from 0 ms to 1000 ms were included in the test (251 total comparisons). When testing time frequency maps, time points from 0ms to 1000ms and all frequency points were included (i.e. 4840 total comparisons). Repeated measures t-tests were performed for each comparison using the original data and 5000 random within-participant permutations of the data. For each permutation, all t-scores corresponding to uncorrected p-values below the threshold of 0.025 were formed into clusters. The sum of the t-scores in each cluster is the “mass” (*t_mass_*) of that cluster. The most extreme cluster mass in each of the 5001 sets of tests was recorded and used to estimate the distribution of the null hypothesis. The family-wise alpha level was set to 0.05. The relatively conservative threshold of 0.025 was chosen to further minimize the likelihood of false positive findings. It is important to note that in the cluster mass permutation test the most extreme *t_mass_* value, likely corresponding to the largest cluster, is used to create a null distribution. Therefore, cluster mass permutation test are useful to capture the broad changes which form large clusters (i.e. span many time- and frequency-points) as these likely have high *t_mass_*. But the cluster mass permutation test might miss focal effects forming clusters of small *t_mass_*, even if specific comparisons within a cluster are highly significant (see: Groppe et al., 2011). We found significant effects mainly in the theta band, but it is possible that significant effects occurred also in other frequencies, but they were not considered significant by the cluster mass permutation procedure because of the high *t_mass_* of the theta-band cluster.

We did not correct for the multiple comparisons across measures, since due to different nature of comparisons (e.g. against baseline, between conditions, with null models) it would not be clear how to define a correction. Yet the reported *p* values were typically low (<0.001), thus additional correction is unlikely to change the study’s results.

The following notation was used to indicate p values in all figures: *p<0.05 **p<0.01 ***p<0.001 ****p<1x10^-7^

### 2.13 Software

The experiment was written and conducted in Matlab Psychotoolbox (Brainard, 1997; Pelli, 1997) running under Matlab 2013. Data analysis was conducted using Matlab 2013 and the following open-access toolboxes: EEGlab (Delorme and Makeig, 2004), Brainstorm (Tadel et al., 2011), Brain Connectivity Toolbox (Rubinov and Sporns, 2010), and Mass Univariate ERP Toolbox (Groppe et al., 2011).

## 3. Results

In the present study we investigated how perception and cognition in the visual oddball task affect brain phase-synchronized networks. The oddball task was employed as a model to study network reorganization as processing of rare ‘oddball’ targets involves several cognitive operations, including reorientation of attention and memory comparisons, and activates widespread brain networks, including fronto-parietal and subcortical areas (Brázdil et al., 2007; Kim, 2014; review: Polich, 2007).

### 3.1 Behavioral results, ERP, and event-related spectral power

The mean detection rate of targets in the visual oddball task was 97±0.009 % and the mean reaction time was 579±16ms. In response to targets ERPs exhibited stronger N1 and N2 peaks (t_mass_=153.6, p=0.0016; **Figure 2A**) and higher amplitude of the P3 peak (t_mass_=376.8, p<0.001) than in response to distractors. Spectral power changes also exhibited the expected pattern (Mazaheri and Picton, 2005). Processing of both distractors and targets led to increase of theta power over baseline (DIST: t_mass_=1861, p<0.001; TARG: t_mass_=2373, p=0.004; **Figure 2B**). Similarly in both conditions alpha/beta desynchronization occurred (DIST: t_mass_=4867, p<0.001; TARG: t=4042, p<0.001) but only after presentation of a distractor there was a beta power increase with onset around 600ms after the stimulus (t_mass_=2162, p<0.001), which is the well-known “beta rebound” phenomenon (Pfurtscheller et al., 2005). Comparisons between conditions indicate that theta synchronization (t_mass_=1658, p=0.006) and alpha desynchronization (t=3648, p<0.001) were stronger during targets processing.

**Fig 2.**
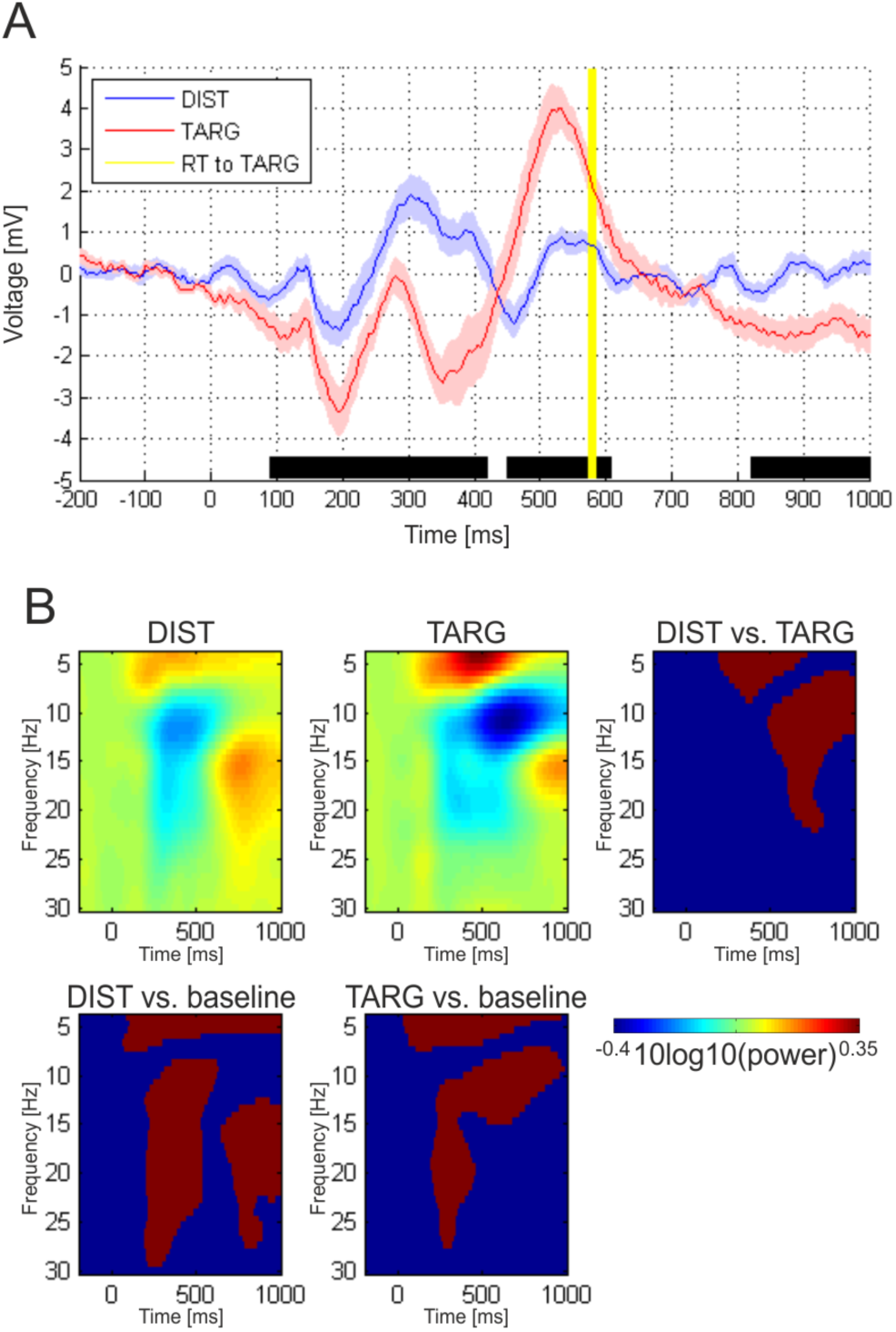
Event-related potentials (ERP) and spectral power in response to distractors (DIST) and targets (TARG) in the oddball task. **(A)** ERPs averaged over 5 parietal electrodes and presented as mean (solid lines) ± SEM (shaded regions). Black bars below ERPs represent time points with significant difference between conditions. Yellow vertical line represents mean reaction time (RT) to targets. **(B)** Spectral power changes averaged over all channels and plotted as 10log10 change over baseline. Two panels below and the panel on the right side show results of the statistical comparisons. Red color indicates time-frequency regions significantly different from baseline (panels below) or between conditions (right side). Blue color indicates no significant difference.

### 3.2 Event-related network analysis (ERNA): network strength

The main aim of the present study was to characterize weighted networks during a cognitive task. Firstly, we analyzed *strength* of interactions within weighted ERNs. *Strength* was calculated as average over weights of all edges in a network. Both, distractors and targets resulted in a network-wide increase of theta band *strength* (DIST: t_mass_=1811, p<0.001 TARG: t_mass_=3245, p<0.001; **Figure 3A, B**) which was stronger during target processing than during distractor processing (t_mass_=2407, p<0.001). Analysis of the *nodal strength* (**Figure 3C**) revealed that initially (around 200ms) only occipital nodes increased strength, but during cognition also parietal and frontal areas are included in the strongly connected task-relevant network. Interestingly, after the manual response only the nodes of the left hemisphere motor areas exhibit higher strength, what would be expected after execution of a right-hand finger movement (e.g. Nolte et al., 2004).

**Fig 3.**
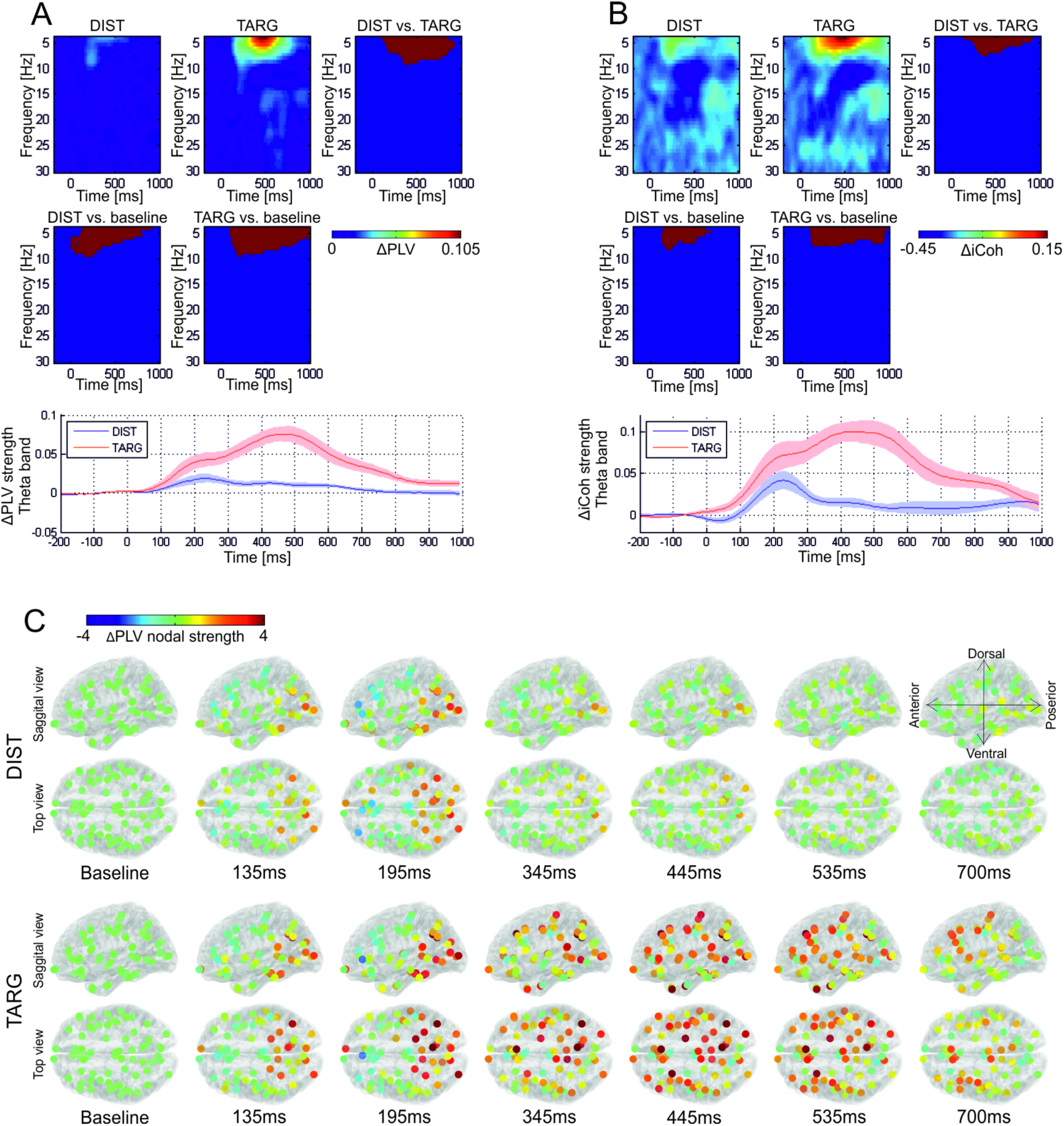
Strength of weighted event-related networks. Results obtained with two connectivity measures are presented: phase locking value (*PLV*; **A, C**) and imaginary part of coherence (i*Coh*; **B**). **(A, B)** Network strength defined as average over weights of all connections is plotted as change over baseline in time-frequency plot (upper panel) and in time for the theta band (lower panel). The panels depict results of statistical comparisons organized as in Fig 2B with red indicating statistically significant difference. **(C)** Change over baseline of *nodal strength* in the theta-band *PLV* network defined as sum of weights of all connections coupled to a node.

To further support our conclusions we created weighted networks using imaginary part of coherence (*iCoh*; Nolte et al., 2004) - a conservative measure of functional connectivity insensitive to volume conduction. Processing of both, targets and distractors, led to an increase of theta-band *iCoh* network *strength* (DIST: t_mass_=1049, p=0.004; TARG: t_mass_=2568, p<0.001). Again, the difference between conditions was apparent, with greater theta *strength* increase during target processing (t_mass_=1524, p<0.001). This provides strong evidence for a network-wide increase of functional coupling during cognitive operations.

### 3.3 Event-related network analysis: clustering and path length

Further, we analyzed topology of the weighted event-related *PLV* networks with a number of graph measures (Rubinov and Sporns, 2010; 2011). C*haracteristic path length* (*CPL*; **Figure 4A**) and *clustering coefficient* (*CC*; **Figure 4B**) were used as the basic measures. We found that perceptual and cognitive processing was related to rapid and transient reorganization of the theta band network’s topology. Specifically, both distractors and targets increased path length (DIST: t_mass_=1461, p=0.0015; TARG: t_mass_=2817, p<0.001) and clustering of the network (DIST: t_mass_=968, p=0.006; TARG: t_mass_=2725, p=0.008). But the increase in both, path length (t_mass_=2290, p<0.001) and clustering (t_mass_=1941, p<0.001), was stronger during cognitive processing of targets. Inspecting the *nodal CC* (**Figure 4C**) we observed an increase of *nodal CC* in a great majority of nodes during cognition which shows that the effect was rather network-wide and not driven by an increase in *CC* in one specific location.

**Fig 4.**
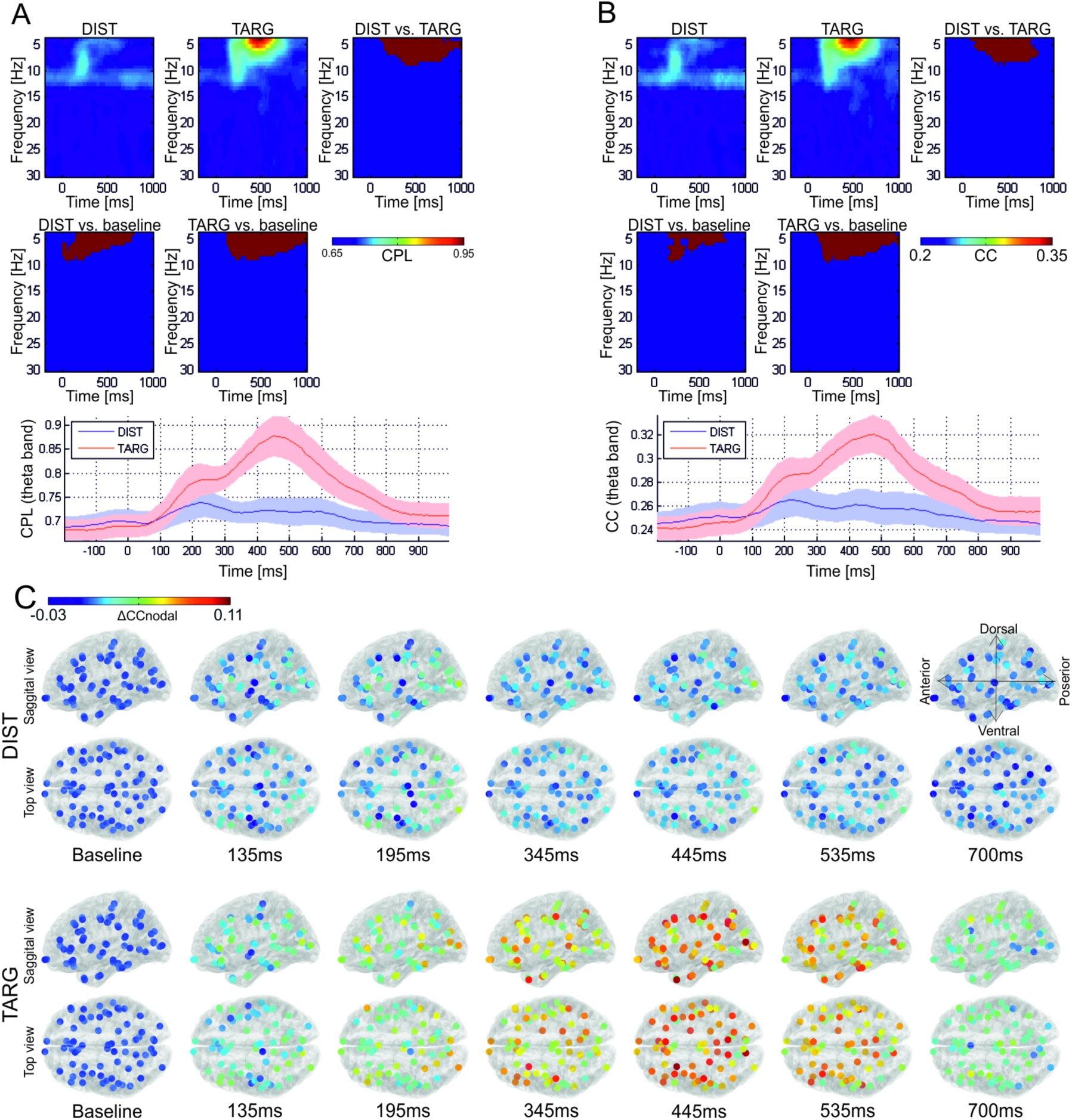
Reorganization of the weighted event-related networks quantified with graph measures. Graph measures plotted in time-frequency plots (upper panels) and in time for the theta band (lower panels). Panels depicting results of statistical comparisons organized in the same manner as in Fig 2 and 3. **(C)** Change over baseline in theta-band *nodal CC*.

### 3.4 Event-related network analysis: modularity

Further, we analyzed dynamic modularity of the event-related networks. A module (community) can be understood as a set of highly inter-connected nodes which are relatively sparsely connected to nodes in other modules. In the present study modularity was studied in two ways (**Figure 5A**). Firstly, we investigated modularity of uni-layer networks where each network represented one time- and frequency-point. Secondly, we studied community structure in the multilayer (temporal) networks where each layer (“slice”) corresponded to one time-point and subsequent layers were linked to each other (Mucha et al., 2010; Bassett et al., 2011, 2013; Doron et al., 2012). In this way we were able to capture not only spatial but also temporal aspects (i.e. temporal evolution) of the community structure.

**Fig. 5.**
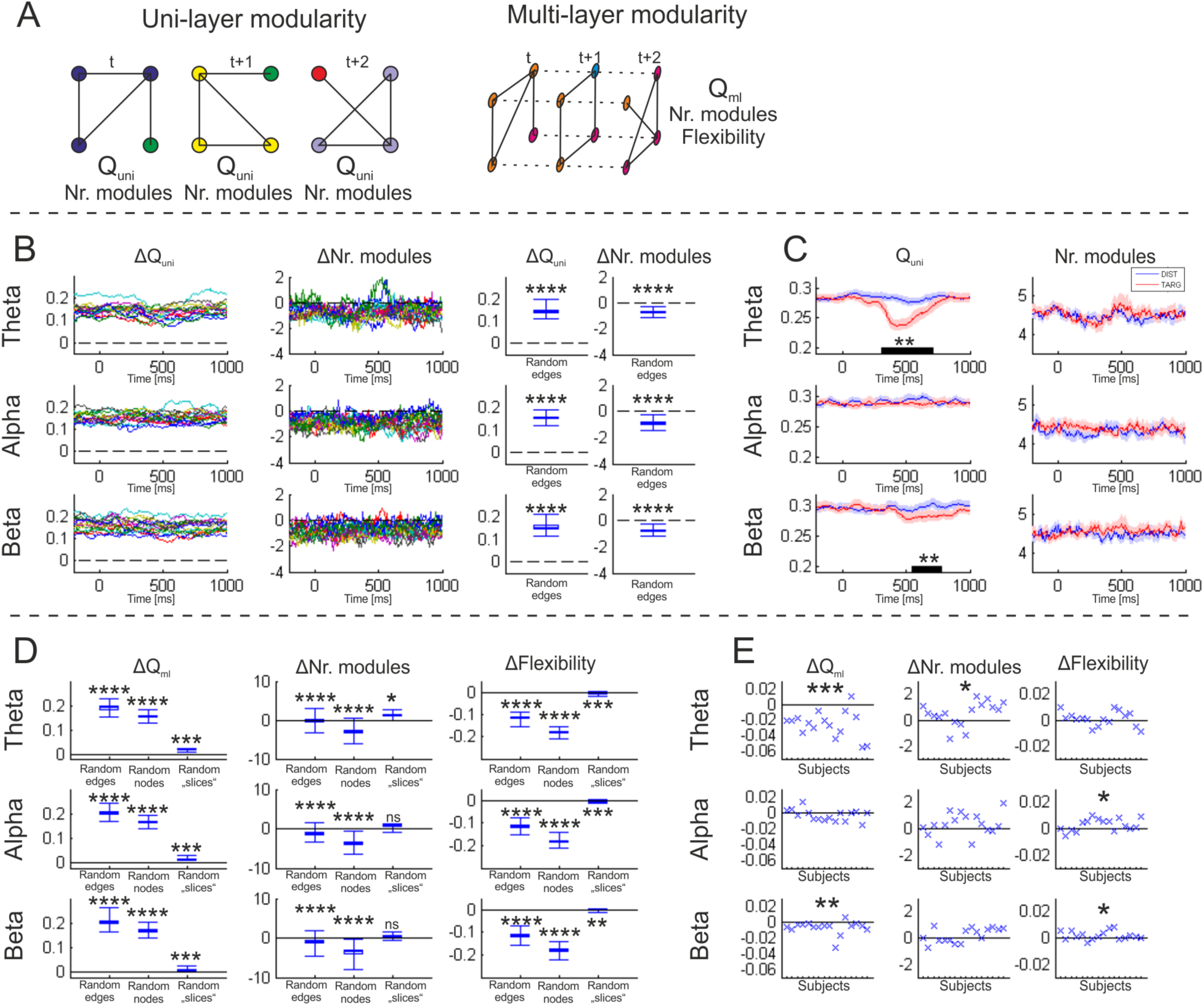
Reorganization of modular structure in the weighted event-related networks. (**A**) Modularity was studied in two types of networks crated from the same data. In uni-layer networks parameters (*Q_uni_* and number of modules) were calculated for each time point separately while in multi-layer networks *Q_ml_*, number of modules, and flexibility were calculated for the whole multilayer network. (**B, D)** The difference (Δ) between features of real uni-layer (**B**) and multilayer (**D**) networks and null networks were calculated for each subject. Both uni-layer and multi-layer networks exhibited modular structure as indicated by higher *Q_uni/ml_* and smaller number of modules than in null models. (**C, E**) Further, to test modularity during cognitive processing network features were compared between conditions. Absolute values are plotted for uni-layer networks and time-points where significant difference between conditions was found are marked with a black bar (**C**). For multi-layer networks the difference between targets and distractors is plotted for each subject (**E**). Result of statistical comparisons: *p<0.05 **p<0.01 ***p<0.001 ****p<1x10^-7^

Yet, analysis and interpretation of the community structure is non-pertinent if a network does not exhibit modular topology. Therefore, we firstly tested whether the analyzed event-related networks are modular by comparing original networks to random null-models. This initial analysis established that both, uni- and multi-layer event-related networks indeed exhibit modular structure. The uni-layer event-related networks were characterized by higher *Q_uni_* and smaller number of modules than random networks (**Figure 5B**). The multi-layer networks exhibited higher *Q_ml_*, smaller number of modules, and lower flexibility when compared to the three null models (**Figure 5D**).

Results of both analyses, uni- and multi-layer networks, indicates that cognitive processing of targets reduces modularity of functional brain networks. The uni-layer theta band network became transiently less modular around 400-500ms after the target presentation, as indicated by lower *Q_uni_* (t_mass_=184, p<0.001; **Figure 5C**). Modularity of the beta band network decreased as well, but later, around 600-700ms after the stimulus onset (t_mass_=85, p=0.0016). Further, analysis of modular structure in the multi-layer networks confirmed that cognitive processing is related to lower modularity of the theta (t(1,15)=5.94, p<0.001) and beta networks (t(1,15)=3.27, p=0.005; **Figure 5E**). In the multi-layer networks we found that during cognitive processing the number of modules in the theta network (t(1,15)=2.18, p=0.045), and flexibility of the alpha (t(1,15)=2.61, p=0.019) and the beta network (t(1,15)=2.31, p=0.034) was higher than during distractor processing. Thus, both analyses demonstrate that cognitive processing is related to changes in the spatio-temporal patterns of networks’ community structure. Specifically, networks topology shifts towards less modular arrangements during cognitive processing which indicates extensive integration of information between local modules.

### 3.5 Event-related network analysis: network hubs

The initial findings - namely an increase of clustering and a decrease of modularity - seemed contradictory at first sight, as more modular networks are typically also more clustered. Yet, we hypothesized that dense connectivity among hub-nodes belonging to different modules might, at the same time, increase nodal clustering of hub-nodes (thus, drive the global clustering increase) and make borders between modules less clear-cut (thus decrease modularity). To test whether cognitive processing is related to greater inter-connectivity between hubs we calculated the Rich Club Coefficient (*RCC*). *RCC* quantifies weights of inter-connections between the rich-club nodes (i.e. nodes with highest *degree*) in relation to general *strength* of the network. During cognitive processing of targets *RCC* increased over baseline in the theta (t_mass_=1793, p=0.0064; **Fig. 6A**) and beta bands (t_mass_=762, p=0.034). In both bands *RCC* increase was in fact stronger after target presentation than after distractor presentation (theta: t_mass_=1601, p=0.0014; beta: t_mass_=404, p=0.045). *RCC* increase was found for a wide range of *rcK* cut-off values, i.e. in rich-clubs comprising more or less nodes (**Fig. 6B**).

**Fig. 6.**
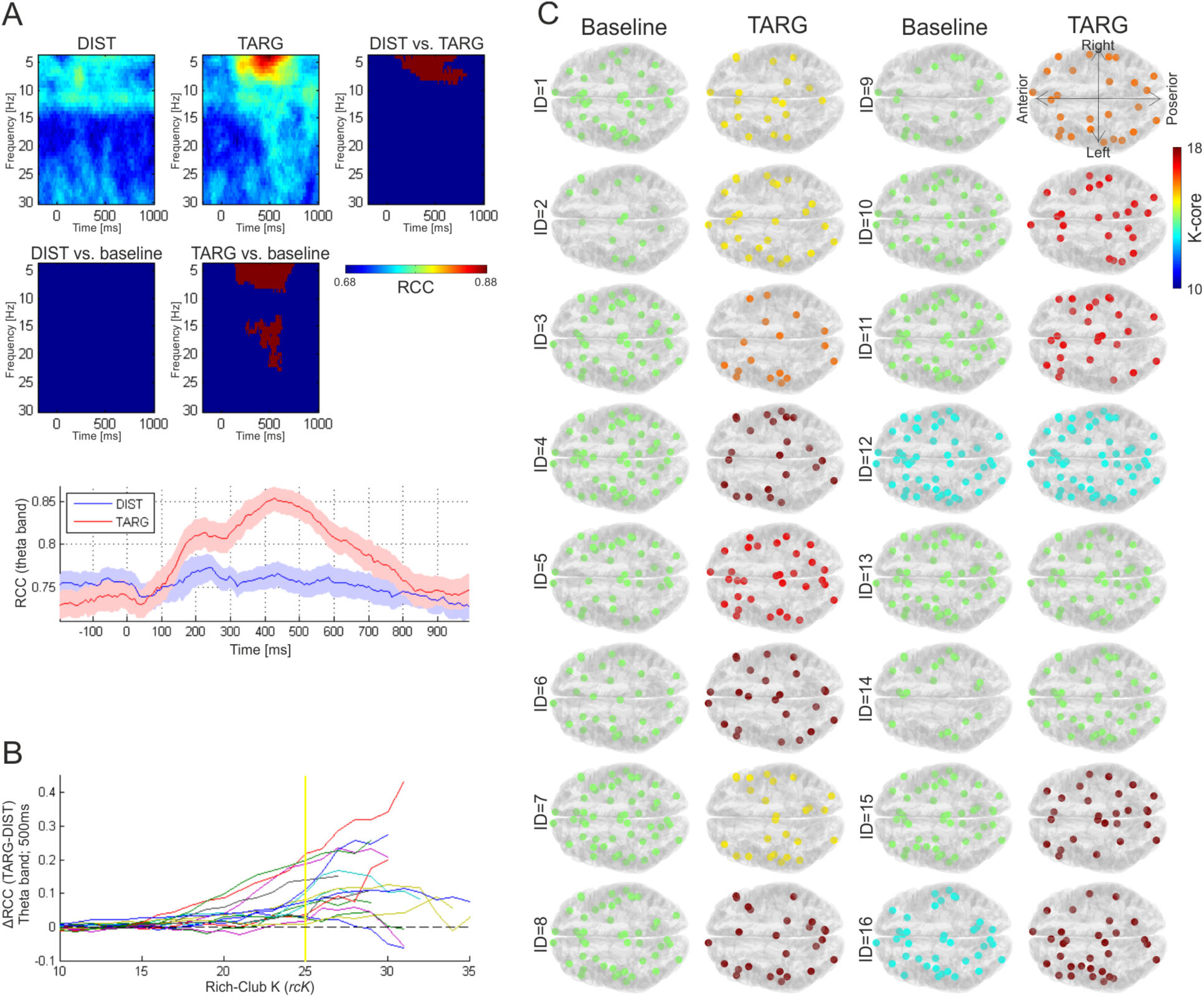
Reorganization of the network core in the event-related networks. (**A**) Rich-Club Coefficient (RCC; *rcK*=25) plotted in the time-frequency plots (upper panel) and in time for the theta band (lower panel). Panels depicting results of statistical comparisons organized in the same manner as in Fig 2 and 3. (**B**) The RCC difference between conditions in the theta band at 500ms plotted as a function of *rcK*. Each subject is plotted in one color and yellow vertical line indicated *rcK* for RCC plotted in **A**. Higher RCC during target processing can be found across wide range of *rcK*. **(C)** K-core of the theta band network at baseline and at 500ms after target presentation plotted for each subject.

Results of the k-core decomposition (**Fig. 6C**) provide further support for the role of extensively connected set of hub-nodes in cognitive processing. The k-core subgraphs found during targets processing were characterized by higher *k-core* values (i.e. included nodes had higher *degree*) and at the same time were sparser (i.e. less nodes were included) than k-core subgraphs found at baseline. Thus a sub-network comprising few interconnected high-degree nodes emerges during cognition. Altogether, our findings support the hypothesis that inter-connectivity among the hub-nodes becomes denser during cognitive processing.

### 3.6 Control analyses

Graph theory has only recently been introduced to neuroscience, thus no well-established preprocessing and analysis pipelines exist. In addition, there are still numerous caveats related to the application of graph measures to neuroscience data (recently reviewed in: Fornito et al., 2013; Papo et al., 2014). When analyzing topology of sparse networks one has to define a number of parameters which might influence the final result. Thus, to demonstrate that the main effects of the study are robust we conducted several control analyses. Specifically, we showed that (i) event-related theta band network reorganization can be found in weighted networks across a wide range of *densities* (**Figure 7A**); (ii) varying network *density*, resolution parameter *gamma*, and coupling parameter *omega* does not change the main results of the community structure analysis (**Figure 7B-F**); and (iii) analysis of binary (un-weighted) networks leads to the same conclusions as analysis of weighted networks (**Figure 8**). Thus, the topological effects cannot be ascribed to the mere increase in network *strength* during stimulus processing.

**Fig. 7.**
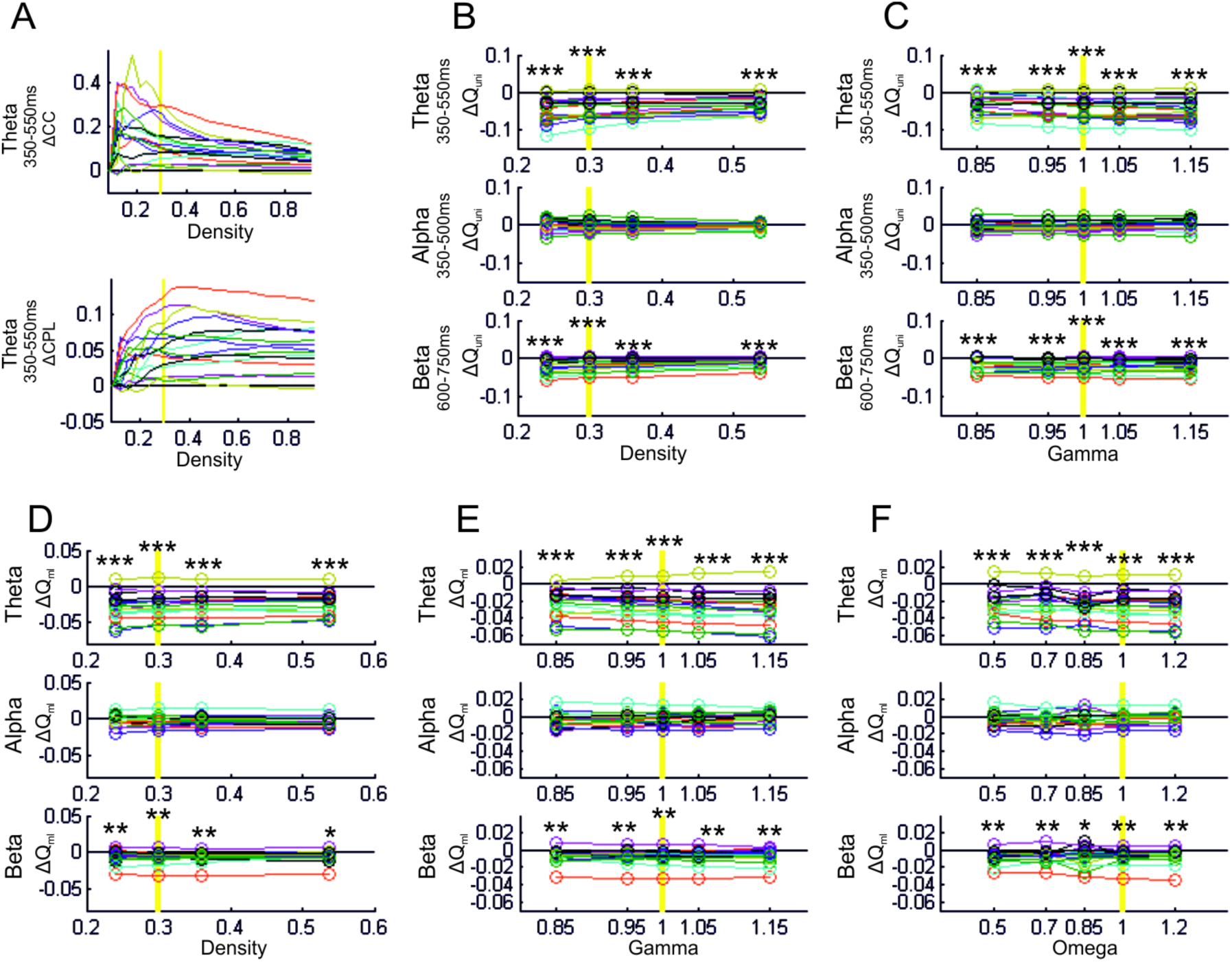
Control analyses of the weighted event-related networks. The yellow vertical line indicates the value of each parameter in the main analysis. In all plots the difference between conditions (TARG-DIST) for each subject (different colors) is depicted. (**A**) *CC* and *CPL* plotted against network density. (**B, C**) Uni-layer modularity was calculated across range of *densities* and resolution parameter *gamma* values. (**D-F**) Multi-layer modularity was calculated across range of densities, resolution parameter *gamma*, and between-layer coupling parameter *omega*. Results of statistical comparisons: *p<0.05 **p<0.01 ***p<0.001 ****p<1x10^-7^

**Fig. 8.**
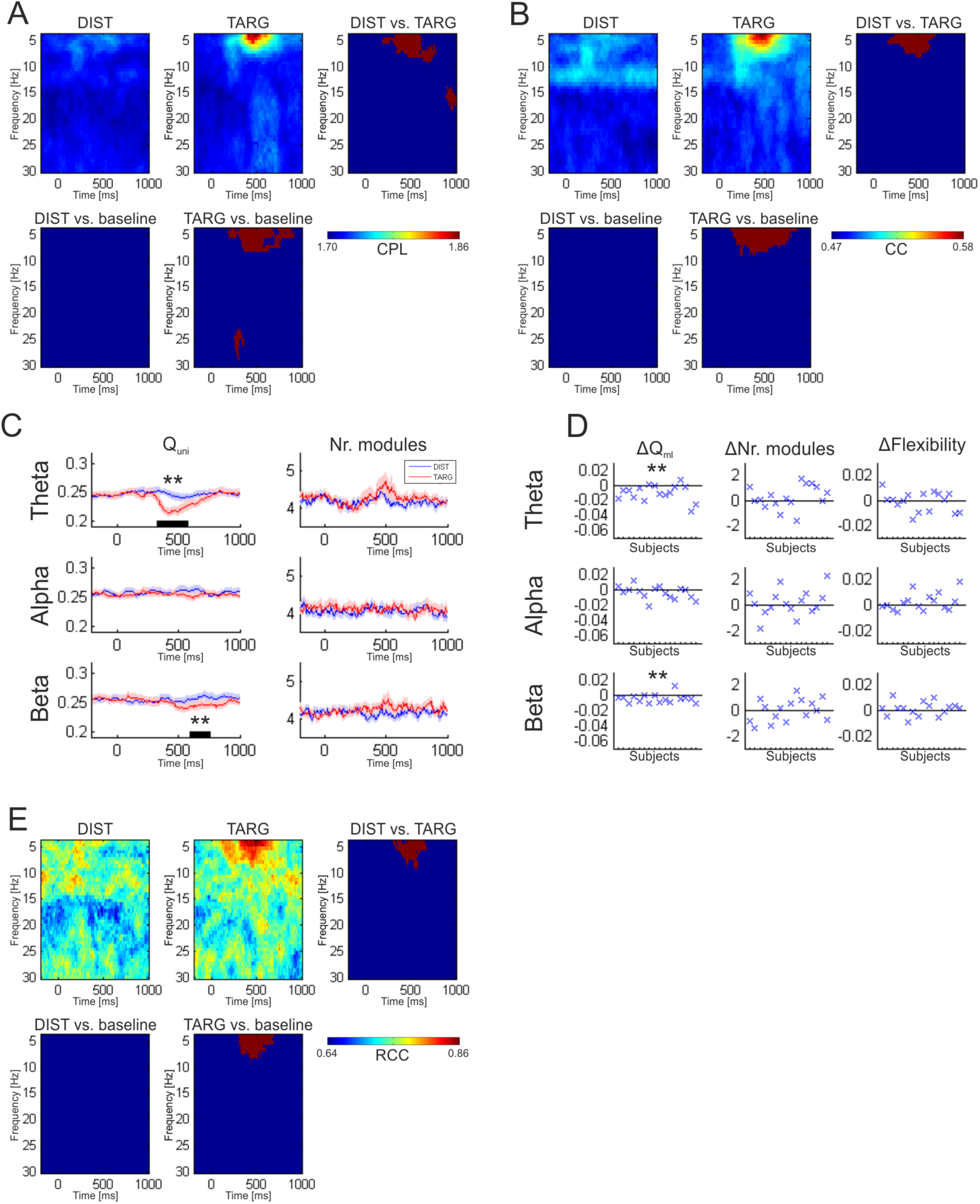
Reorganization of event-related binary networks quantified with graph measures. (**A-B**) Graph measures plotted in time-frequency plots. Panels organized as in Fig. 4. Modularity of uni-layer (**C**) and multi-layer binary networks (**D**) plotted as in Fig. 5. (**E**) Rich-Club Coefficient (RCC). Results of statistical comparisons: *p<0.05 **p<0.01 ***p<0.001 ****p<1x10^-7^

## 4. Discussion

Cognition and consciousness are believed to emerge from activity of widespread brain areas coordinated by phase synchronization (Gaillard et al., 2009; Hipp et al., 2011; review: Varela et al., 2001; Siegel et al., 2012). Such interactions are highly dynamic with functional networks established and dissolved on the time scale of tens of milliseconds. Yet, not much is known about the topological arrangements of transient cognitive networks, as graph theory has been employed predominantly to study time-invariant anatomical and functional-resting-state networks. Here we used a visual oddball task as a model of perceptual and cognitive processing (review: Polich, 2007). Subjects were asked to ignore frequent distractor stimuli and react with a button press to rare targets only. Target recognition involves several cognitive operations, e.g. reorienting of attention and memory comparisons, and activates fronto-parietal brain networks (Brázdil et al., 2007; Kim, 2014; review: Polich, 2007). Using the excellent temporal resolution of EEG we revealed that cognitive processing is related to rapid, transient, and frequency-specific reorganization of functional networks’ topology as demonstrated by our event-related network analysis (ERNA). Specifically, cognitive networks were characterized by three features: (i) strong clustering, (ii) low modularity, and (iii) strong interactions between the hub-nodes. Our findings suggest that dense and highly clustered connectivity among hub-nodes belonging to different modules is established during cognition. Such inter-modular connections might be a substrate of the “global workspace” necessary for the emergence of conscious perception (Baars, 2002; Baars et al., 2013; Dehaene and Changeux, 2011).

### 4.1 Global topology of brain networks: fixed or flexible?

The majority of hitherto conducted studies on brain functional connectivity aimed to map the spatial profile of spontaneous brain networks by assuming their temporal invariance. Yet, recently, it was demonstrated that the functional networks are highly dynamic and non-stationary (MEG/EEG: Betzel, et al. 2012; Baker et al. 2014; fMRI: Allen et al., 2014; Zalesky et al., 2014). Such dynamics is related to changes of psychophysiological states on a slow-time scale of minutes (Chang et al., 2013). Nevertheless, it is not clear whether topology of intrinsic brain networks reorganizes on a fast sub-second time-scale. It is conceivable that reorganizing into qualitatively new, more efficient topological patterns might support cognitive processing. Yet, the hitherto conducted studies on task-related network reorganization were sparse and inconclusive. A number of reports concluded that cognitive processes merely change weights or local (nodal) features of intrinsic functional networks but not the global topological structure (Bassett et al., 2006; Nicol et al., 2012; Betti et al., 2012; Jin et al., 2012; Crossley et al., 2013; Cole et al., 2014). Yet, others observed task-related topological reorganization of large-scale networks on a time-scale of seconds (Valencia et al., 2008; Doron et al., 2012; Palva et al., 2010; Kitzbichler et al., 2011; de Vico Fallani et al., 2008; Ekman et al., 2012). On a longer time scale of minutes/hours, fMRI network’s rearrangements are related to plasticity, e.g. motor learning (Bassett et al., 2011; 2013).

Results of the present study reveal that reorganization of the network’s *topology* might contribute to cognitive processing, beyond the mere increase in *strength* of functional coupling. The key role of brain functional networks’ topology in cognition has already been postulated by the global workspace theory (GWT; Baars, 2002). The assumption of GWT is that after local processing within specific modules (e.g. visual, auditory, motor systems) the information needs to be integrated within a global workspace, which can be identified with a network comprising hub-nodes and inter-modular connections. Hence, according to GWT, it is not the strength of pairwise connections but rather the topological arrangement of a network which gives rise to perception, cognition, and consciousness. The reorganization pattern observed in the present study, namely a decrease of modularity and strong interactions between hubs, is in good agreement with the predictions of GWT. Interestingly, a decrease of modularity and shift towards global integration were also observed in the previous studies analyzing task-related network reorganization (Kitzbichler et al., 2011; Kinnison et al., 2012; Ekman et al., 2012). Thus, the application of graph theory to neuroimaging data provides a growing body of evidence in favor of the GWT.

When interpreting results of this and previous studies and pondering the aforementioned discrepancies and similarities it is important to keep in mind that functional connectivity is a broad concept including various types of interactions (Friston, 2011). Recently, it was proposed that two different functional coupling/connectivity modes exist: (i) slow envelope coupling captured by correlations between BOLD signals and (ii) dynamic phase coupling captured by phase locking (Engel et al., 2013; Mehrkanoon et al., 2014). While the envelope coupling has a “modulatory” influence on cognition, the phase coupling is identified with cognitive processing itself. Therefore, the networks representing phase coupling are more likely to dynamically adjust topology during cognition, which is what we observed. Yet, it is important to note that also fMRI studies found task-related changes in global topology (Ekman et al., 2012; Bassett et al., 2011) hence the relationship between the coupling modes and the task-related reorganization might be far from obvious.

The modules studied in the event-related EEG networks (both uni- and multi-layer) might be interpreted as functionally coherent entities. Notably, the modules we observed were not *spatially* coherent, i.e. distant nodes often belong to the same module while adjacent nodes belong to different modules (see: **Figure 9**). This is in sharp contrast to previous studies of anatomical networks and fMRI functional networks where spatially coherent modules were found (review: Meunier et al., 2010). Yet, this discrepancy might in fact be explained by the two aforementioned modes of coupling. Actually, the postulated mechanism of cognition is transient phase-synchronization of “neural coalitions” comprising *spatially distributed* neural assemblies (Crick and Koch, 2003), and in the graph theory framework such “coalitions” might be identified with spatially widespread, incoherent modules.

**Fig. 9.**
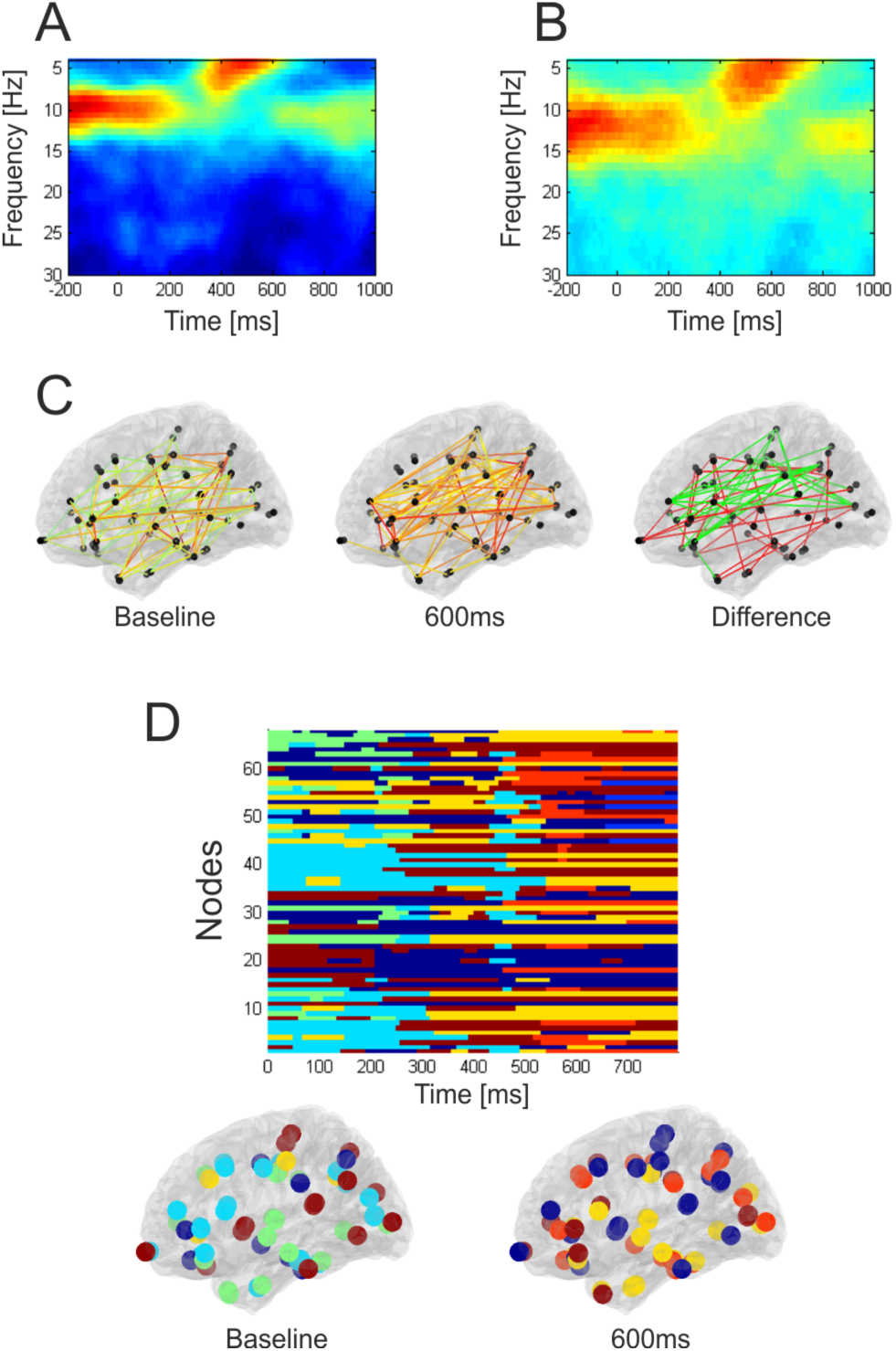
Weighted event-related networks of cognitive processing: a single-subject analysis. (**A**) Network strength and (**B**) clustering coefficient plotted in time-frequency plots. (**C**) Snapshots of the theta-band weighted network (*density*=0.12) at two time-points. The difference plot depicts edges present at baseline but missing during cognitive processing (in red), and edges present during cognitive processing but missing at baseline (in green). (**D**) Spatio-temporal dynamics of community structure in the theta band network (multi-layer analysis). Each module is represented by one color. Below, snapshots of the community structure at two time-points.

### 4.2 The architecture of cognitive networks

We revealed reorganization of functional large-scale networks’ during cognition but two questions remain. Firstly, what is the exact pattern of network reorganization, and secondly, what is the functional role of the reorganization pattern in cognition (i.e. how can it support information processing). A wide range of graph measures used in our study provided a comprehensive picture of the cognition-related reorganization and thus allowed us to propose a specific “fingerprint” of event-related reorganization.

Network clustering is typically strong in modular networks where modules comprise highly interconnected nodes. Thus, the two simultaneous occurring effects - increase of clustering and decrease of modularity - seem to be at odds with each other. Yet, it is the case only if one assumes that the increase of clustering must take place *within* modules. Conversely, we hypothesized that an increase in clustering is driven by connections established between modules, i.e. between the hub-nodes. If this is the case, then such inter-modular connections cause also decrease of modularity, as the initially clear-cut borders between modules become vague (**Fig. 10**). The analysis of *RCC* seems to confirm this hypothesis as inter-connectivity between rich-club nodes increased during cognition. Therefore, although increased clustering is typically interpreted as a shift towards local segregation of information within modules, here we view increasing clustering as indicative of dense, clustered connectivity between hub-nodes which allows extensive global integration of information (Sporns, 2013). Importantly, previous studies investigating cognitive networks across a range of tasks and using different analysis methods found a similar decrease of modularity and shift towards global integration of information (Kitzbichler et al., 2011; Kinnison et al., 2012; Ekman et al., 2012). This raises the possibility that this particular reorganization pattern accompanies different cognitive processes and is a genuine network “fingerprint” of cognition. This hypothesis needs to be tested by future studies.

**Fig. 10.**
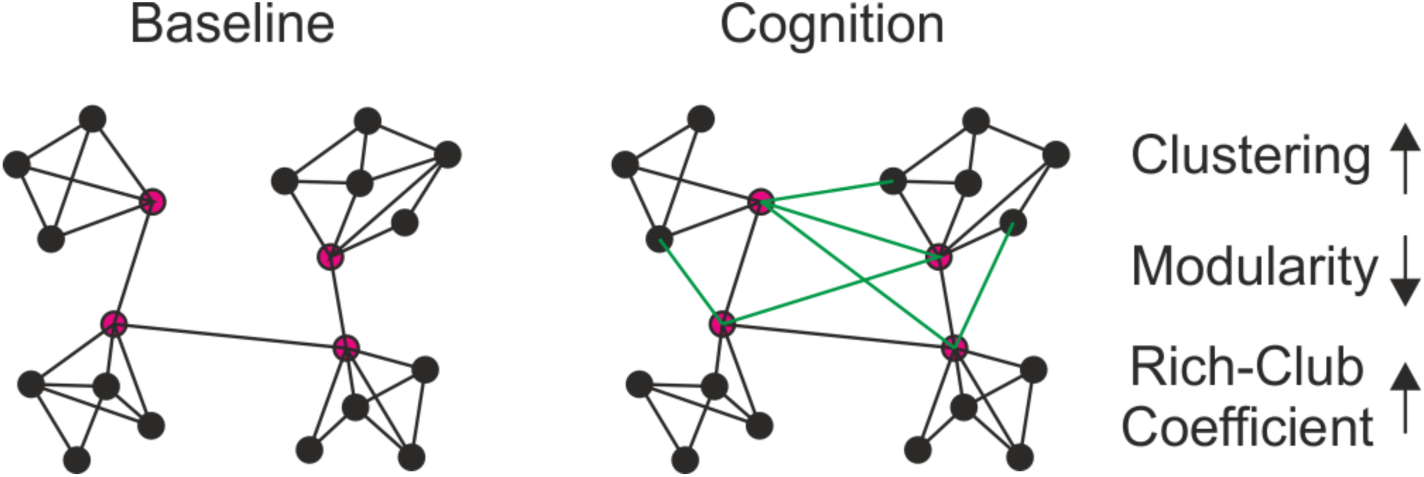
The proposed network topological “fingerprint” of cognition. In this scheme baseline network is characterized by modular structure (with four modules here) and high clustering within modules, in agreement with the literature. Hub-nodes (in magenta) exhibit high *degree* and transfer information across modules. We propose that dense and clustered inter-modular connectivity (in green) among hub-nodes is established during cognition. Such reorganization pattern accounts for the three findings of the present study, namely: (i) increase in clustering; (ii) decrease in modularity; (iii) increase in Rich-Club Coefficient.

The hypothesis that inter-modular connections are dynamically established between hub-nodes is in line with recent fMRI studies. The most dynamic connections in the resting-state are indeed the inter-modular connections (Zalesky et al. 2014) and flexibility of the inter-modular connectivity of hub-nodes is crucial for task execution (Cole et al. 2013, 2014) and for the self-generated thoughts (Schaefer et al. 2014). The intermodular connections are also most costly in terms of energy (Bullmore and Sporns, 2012), and thus might be established primarily during execution of a demanding task, but to a lesser extent at rest. Importantly, disruption of the hub-nodes is believed to be the common mechanism of neurological and neuropsychiatric disorders affecting cognition (Crossley et al., 2014).

Changes in clustering of event-related networks might also inform us whether cognitive networks exhibit rather “hierarchical” or rather “parallel” architecture. In the hierarchical architecture one hub-area is essential for processing and all other brain areas involved are coupled to the hub but not to each other. Conversely, in a flexible and parallel architecture several regions might exchange information but none of them plays the central or “lead” role. Hierarchical and parallel architecture would be characterized by low or high clustering coefficient, respectively. Thus, the observed increase in clustering lends support to the model of parallel architecture of the task-related networks. The strong k-cores provide further evidence for strong, parallel sub-network linking nodes most important to the task. Interestingly, a parallel, small-world architecture with strong interactions between core and periphery present at the stimulus onset facilitates stimulus processing and task performance (Ekman et al. 2012; Weisz et al., 2014). Yet, it is also important to consider the transitivity of correlation measures (i.e. if A is correlated with B, and A is correlated with C, then B is likely correlated with C) which is one of the limitations of functional connectivity analysis (see: Zalesky et al., 2012; Fornito et al., 2013). Due to this property functional networks are by definition clustered, and that might lead to overestimation of the “parallel” network circuits.

Graph theory has been extensively applied in clinical neuroscience (e.g. Bola et al., 2014; Lord et al., 2012), but the great majority of studies investigated the resting-state networks. The resting-state paradigm has several advantages (e.g. patients who are not able to perform a task might be enrolled in the study) and the spontaneous activity is intimately related to the task-evoked activation patterns (review: Sadaghiani et al., 2010). Yet, it would be of importance to test whether the event-related networks studied here can provide deeper insight into mechanisms of various pathological conditions affecting perception and cognition. Possibly, cognitive dysfunctions at the early stage might not be manifested in the resting-state networks, but only in the event-related networks, i.e. when subjects are performing a demanding task. This hypothesis can be tested by future studies.

### 4.3 Methodological considerations and limitations

In the present study we employed the visual oddball task to study networks reorganization. The oddball task is a well-known model to study perceptual and cognitive processing, often employed in the clinical setting. Another reason to choose this task was that processing and reaction to rare ‘oddball’ targets involve numerous basic cognitive operations (e.g. reorientation of attention, memory operations) and activates widespread brain networks (Brázdil et al., 2007; Kim, 2014; review: Polich, 2007). Involvement of several network’s nodes in the task performance makes it also a good model to characterize network’s topological reorganization. Yet, at the same time, using the oddball task might be considered a disadvantage, since “cognition” is then defined in a broad and unspecific manner. Thus, further studies are needed to reveal network correlates of more specific cognitive processes.

Further, we did not find correlations between performance (i.e. reaction time, RT) and network measures. Yet, the oddball task was very easy for the subjects and the variability of RT, both within and between subjects, was low. We hypothesize that the event-related networks measures might be related to behavior when more demanding tasks are used. Such relation between network organization and behavioral measures was indeed observed in previous studies (Ekam et al., 2012; Weisz et al., 2014).

When analyzing large-scale neurophysiological networks using MEG/EEG the common problem is nonphysiological spread of electrical activity through volume conduction causing spurious correlations between signals. Here, the wMNE algorithm was used to address this problem. wMNE is believed to be an optimal source reconstruction methods for large-scale functional connectivity analysis (Hassan et al., 2014; Palva and Palva, 2012). Yet, it is important to keep in mind that the source reconstruction algorithms can only reduce the volume conduction problem, not address it completely.

Another issue to consider is that common feed-forward input and stimulus-locked transients might cause phase-reset in a number of areas, which will artificially increase phase-synchronization among them despite absence of genuine functional interactions. This problem actually occurs in all studies investigating event-related connectivity, and even in the resting-state input from area A might simultaneously phase-reset areas B and C resulting in detection of functional connectivity between B and C without any true interaction. Yet, the key contrast in the present study was between conditions (i.e. DIST vs. TARG) and the stimulus-related transients were present in both conditions making it unlikely that they can account for the observed between-conditions differences. Furthermore, our analysis was focused on the “cognitive” period (300-600ms after stimulus onset) and activity within this time-window is rather not caused by the initial feed-forward sweep of activity. One of the proposed methods to address the stimulus-transient problem is to subtract the average stimulus-locked response (i.e. ERP) from each single trial. But we did not use this method here as such ERP-subtracted signals exhibit unknown properties, particularly in the “cognitive” time-window where brain responses exhibit significant latency- and amplitude-variability (Truccolo et al., 2002). Therefore, developing new (possibly multivariate) analysis techniques more reliably detecting genuine interactions is required to ultimately address this problem.

Finally, although the majority of neuroscience studies employed binary (un-weighted) graphs, here we focused on the weighted graphs which contain more information and thus might ensure greater sensitivity (e.g. Rubinov et al., 2009). Indeed, weighted networks seem to be more sensitive as they indicate topological changes also in response to distractors (**Fig. 4**), while binary networks did not capture this effect (**Fig. 8**). We also noticed that results of the analysis of weighted networks depend to a lesser extent on chosen parameters (e.g. density of the networks). But importantly, analyses of weighted and binary graph were in good agreement with respect to the main effects of the study (i.e. theta-band network reorganization).

### 4.4 Conclusions

The brain functional networks are highly dynamic and able to adjust topology on a very fine time-scale. Here we propose that dense and clustered connectivity between hub-nodes belonging to different modules is the network “fingerprint” of cognition. Such rearrangement might support global integration of information among specific subsystems and provide network substrate for the global workspace (Baars, 2002). It remains to be studied whether the same “network fingerprint” is common to all cognitive operations and whether its’ disruption can be observed in pathological states where cognitive processing is impaired.

**Conflict of interest:** The authors declare no competing financial interests.

## Acknowledgements

The study was supported by Otto-von-Guericke University of Magdeburg, and by the BMBF network ERA-net Neuron “Restoration of Vision after Stroke (REVIS)” (Grant nr 01EW1210).

